# Improved methods for bulk cultivation and fixation of *Loxodes* ciliates for fluorescence microscopy

**DOI:** 10.1101/2022.05.29.493905

**Authors:** Brandon Kwee Boon Seah, Christiane Emmerich, Aditi Singh, Estienne Swart

**Author notes:** Corresponding author: Brandon Kwee Boon Seah.

## Abstract

*Loxodes* is one of the best ecologically characterized ciliate genera with numerous intriguing physiological abilities, including gravity-sensing organelles and nitrate respiration. However, these cells have been considered challenging to cultivate in bulk, and are poorly preserved by conventional fixatives used for fluorescence microscopy. Here we describe methods to grow and harvest *Loxodes* cells in bulk with liquid soil extract medium, as well as a new fixative called ZFAE (zinc sulfate, formaldehyde, acetic acid, ethanol) that can fix *Loxodes* cells more effectively than buffered formaldehyde or methanol. We show that ZFAE is compatible with immunofluorescence and the nuclear stain DAPI. *Loxodes* is thus now amenable to long-term maintenance, large-scale growth, and modern cell biology investigations of monoclonal strains in laboratory conditions.

## Introduction

The ciliate genus *Loxodes* was the subject of pioneering ecophysiological studies by Bland Finlay and his colleagues, who used both natural populations and laboratory cultures in their work. *Loxodes* are microaerophiles that can migrate vertically in response to oxygen gradients in the water column, orienting themselves with the help of gravity-sensing organelles called Müller vesicles (Fenchel and Finlay, 1986; Finlay, 1981; Finlay et al., 1986; Goulder, 1980). They are also able to respire nitrate, a trait that is unusual for eukaryotes and which gives them an advantage under anoxic conditions (Finlay et al., 1983).

Despite their intriguing biology, *Loxodes* can be challenging to work with in the laboratory. Although they are sometimes referred to as “uncultivable”, several methods for maintaining cultures have previously been reported (Bobyleva, 1980; Buonanno et al., 2005; Nagel et al., 1997; Neugebauer et al., 1998). A strain of *L. striatus* was available from the CCAP culture collection in the 1990s (Hemmersbach et al., 1996), but appears to have been lost. Most cultivation approaches are similar to that used by (Fenchel and Finlay, 1984), namely growing them in test tubes filled with liquid medium and a layer of soil at the bottom. Respiration by bacteria produces an oxygen gradient in the tube that mimics the natural habitats of these ciliates (Finlay et al., 1986). However, this format with solid soil particles is inconvenient for growing and harvesting large quantities of cells for experiments.

Furthermore, conventional fixatives used in histology and molecular biology, such as buffered formaldehyde solutions or alcohols, perform poorly in preserving the morphology of *Loxodes* and members of the class Karyorelictea in general (Foissner, 2014; Munyenyembe et al., 2021). “Strong” fixatives that are good at preserving ciliate morphology, on the other hand, usually contain toxic and hazardous substances, e.g. Parducz’s solution (mercuric chloride and osmium tetroxide), Stieve’s solution (mercuric chloride), and Bouin’s solution (picric acid). Such fixatives are still used in standard protocols to fix ciliates for morphological description, despite their hazards and the additional costs associated with proper handling and disposal. They are typically used to prepare samples for electron microscopy or silver staining methods (e.g. protargol or silver nitrate) that reveal cytoskeletal structures important to ciliate taxonomy, but may be incompatible with other desired applications, such as fluorescent labels or immunocytochemistry. For example, fixation with osmium tetroxide produces osmium precipitates that interfere with the detection of fluorescent signals (Dyal et al., 1995).

Here, we describe protocols used in our laboratory for growing *Loxodes* cells in bulk using liquid media, as well as a new fixative that we call ZFAE (zinc sulfate, formaldehyde, acetic acid, ethanol) which keeps *Loxodes* cells intact and which is compatible with fluorescence microscopy while avoiding mercuric chloride and other hazardous substances.

## Methods

### Ciliate strains

*Loxodes magnus* strain Lm5 and *L. striatus* strain Lb1 were isolated from local ponds in Tübingen, Germany and Bern, Switzerland respectively by manual pipetting of single cells into soil-liquid tubes. Species identity was determined by nuclei number and morphology with reference to (Wang et al., 2019).

*Blepharisma stoltei* strain ATCC 30299 was obtained from the lab of T. Harumoto (Nara Women’s University, Japan), and grown in SMB-III medium (Miyake, 1981).

### Media for algal culture

Algae for feeding the ciliates were grown in Tris-acetate-phosphate (TAP) medium (10 mL 5× Beijerinck’s solution, 1 mL phosphate buffer, 2 M Tris base, 1 mL Hutner’s trace element solution, 1 mL glacial acetic acid, dissolved in 1 L water) or Tris-phosphate (TP) medium (same as TAP, except acetic acid omitted and pH adjusted to 7.0 with hydrochloric acid).

5× Beijerinck’s solution: 20 g ammonium chloride, 5 g magnesium sulfate heptahydrate, 2.5 g calcium chloride dihydrate in 500 mL total volume of deionized water. Calcium chloride was dissolved separately from the other salts; both solutions were then combined and adjusted to the final volume.

Phosphate buffer: 10.8 g anhydrous potassium hydrogen phosphate and 5.6 g anhydrous potassium dihydrogenphosphate dissolved in 100 mL water.

Hutner’s trace element solution: 50.0 g EDTA disodium, 11.14 g boric acid, 22.0 g zinc sulfate heptahydrate, 5.1 g manganese (II) chloride tetrahydrate, 5.0 g iron (II) sulfate heptahydrate, 1.6 g cobalt (II) chloride hexahydrate, 1.6 g copper (II) sulfate pentahydrate, and 1.1 g ammonium heptamolybdate tetrahydrate in 1 L deionized water. Components except EDTA were dissolved in 550 mL water at 70 °C, EDTA was separately added to 250 mL water and heated until dissolved. EDTA solution was added to the other salts, and the mixture was heated to boiling, then cooled to 70-75 °C. pH was adjusted to 6.5-6.8 with 20% (w/v) potassium hydroxide while temperature was above 70 °C. Solution was diluted to 1 L final volume with deionized water. Container was loosely covered with a cotton plug, and stirred for two weeks at room temperature until color changed from green to purple. Solution was filtered and stored at 4 °C.

### Cultivation and harvesting of algae

Stock solutions of *Chlamydomonas reinhardtii* strain SAG 33.89 (obtained from the Culture Collection of Algae, SAG, University of Göttingen) were grown in 50 mL TP medium, subcultured every two weeks (1 mL inoculum). Bulk cultures for feeding ciliates were grown in 50 to 400 mL TAP medium, inoculated 3-4 days before harvest with 125 µL inoculum from saturated stock culture per 100 mL final volume. Cultures were maintained in Erlenmeyer flasks with loosely fitting aluminum caps under commercially available aquarium LED lamps.

Algal density was monitored by optical density (OD) at 550 and 750 nm with a spectrophotometer (BioRad SmartSpec Plus). OD of saturated algae was about 1.5 for both wavelengths. If cultures were too dilute or concentrated the resuspension volume was adjusted accordingly. Algae were concentrated by centrifugation (1000 rcf; 3 min; room temperature), supernatant was discarded, and the pellet then resuspended either in Volvic water or SMB medium at 1/10 the original volume. Concentrated resuspended algae was used to feed the ciliates.

### Maintenance of Loxodes stock cultures in soil-liquid test tubes

Loamy garden soil was air-dried in open trays and then stored in plastic buckets at room temperature. Soil was sieved through a stainless steel sieve (2.2 × 2.2 mm) before use, to break up clumps and remove stones and large organic debris. 1 to 1.5 g sieved dried soil was portioned into 30 mL glass test tubes with loose-fitting aluminum caps. Soil-filled tubes were autoclaved (121ºC, 20 min), allowed to cool overnight, and autoclaved again, to kill potential eukaryotic spores and cysts. Autoclaved (121 ºC, 15 min), cooled Volvic mineral water was added to test tubes up to the 25 mL mark. Tubes were briefly vortexed and tapped to dislodge air bubbles in the soil, then left for at least two days to allow fine particles to settle out before use. Tubes have been prepared up to three months in advance. An alternative method of autoclaving tubes already filled with both soil and water (Bobyleva, 1980) was less successful for us, because the mixture was often ejected from the tubes during autoclaving.

To inoculate new cultures, 100 to 500 µL of a dense culture (Figure 1A, B) was transferred by pipette (glass or disposable plastic) to a new test tube. Cultures were kept out of direct light in a closed cupboard at stable room temperature (around 24 ºC in our laboratory), and fed once weekly with 50 µL concentrated *Chlamydomonas* per test tube. Cultures were subcultured once monthly using one- or two-month old cultures.

**Figure 1.**
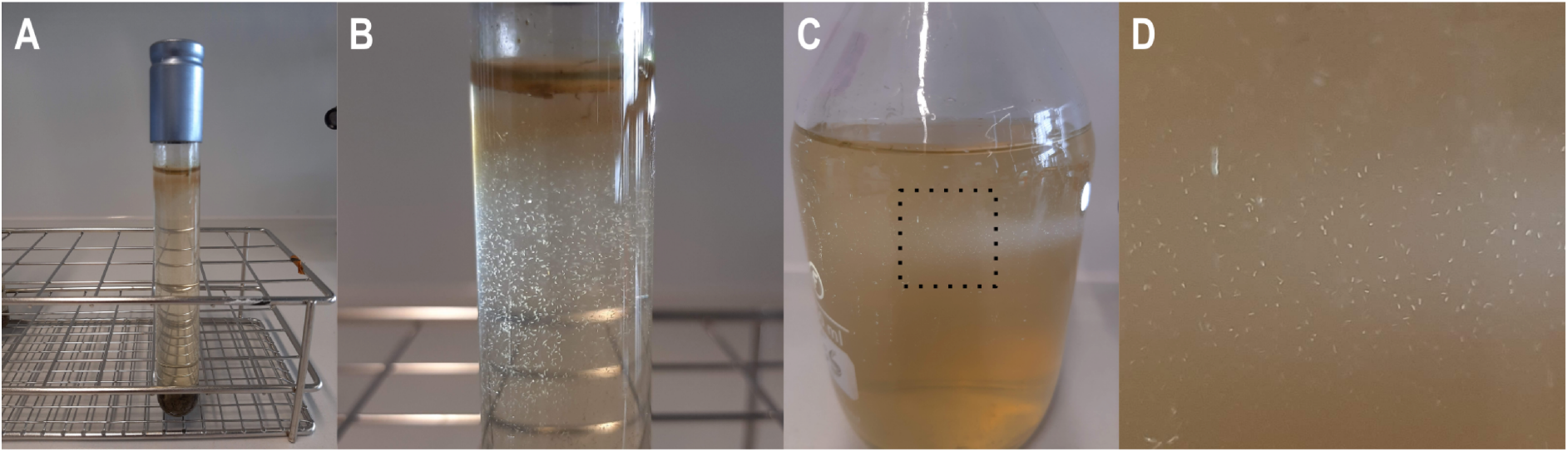
*Loxodes magnus* in soil-liquid test tube (A, B), and soil extract medium (C, D). Close-ups with raking light showing cells in medium (B, D). The dark brown ring on the inner test tube wall below the meniscus is a biofilm produced by ambient microbes.

### Bulk culture of Loxodes in liquid soil extract medium

To prepare soil extract, 82 g of sieved dried garden soil (above) and 600 ml tap water were mixed in a 1 l borosilicate glass bottle, then autoclaved (121 °C, 15 min). After cooling to room temperature and allowing particles to settle, the clear supernatant was decanted into a new bottle, diluted 1:1 (v/v) with tap water, and autoclaved again. Soil extract was stored at room temperature until use. Measurements of total organic carbon, inorganic ions, and elemental composition were performed by SGS lnstitut Fresenius (Taunusstein, Germany); data are available at doi:10.17617/3.4ZQNBK.

To inoculate a soil extract medium culture, >1 ml of a dense stock culture was used to inoculate 50 ml of soil extract medium in a borosilicate glass bottle, taking care not to transfer soil particles. Bottles were closed but still had air-filled headspace in the conical part of the bottle (Figure 1C, D). Cultures were transferred to larger bottles and diluted 1:1 with new soil extract medium when dense, until the desired volume and cell density were achieved. Starting with a smaller volume of soil extract and doubling it when dense was usually more successful than immediately inoculating a large volume (e.g. 500 ml) with a small stock inoculum.

Cultures were fed twice weekly with 300 µl concentrated *Chlamydomonas* per 100 ml culture volume.

### Monitoring of individual cells in glass wells kept in a gas chamber

Clonal lines can also be grown from single cells isolated in three-well glass depression slides, similar to protocols for *Paramecium* (Beisson et al., 2010a; Sonneborn, 1950). Each well was filled with 250 µl soil extract medium, initially fed with 1-5 µl concentrated *Chlamydomonas*, and maintained for several days in polycarbonate gas jars (A05077, Don Whitley Scientific) under microaerobic conditions using N_2_ gas (Whitley Jar Gassing System, Don Whitley Scientific). The bottom of each jar was lined with moist paper towels to prevent evaporation of media from the wells. Three-well slides were placed in plastic Petri dishes to allow them to be stacked in the jar.

### Cell counting and calculation of growth rates

Bulk liquid cultures were gently swirled to distribute cells more evenly before sampling. For counting, three 100 µL aliquots were taken with a wide-bore pipette tip, and gently spotted as a row of smaller droplets on a plastic Petri dish to facilitate counting. Cells were counted under a binocular microscope with the help of a clicker. If density was too high (above about 150 cells / 100 µL), samples were first diluted 10× in SMB or soil extract medium before spotting onto a Petri dish.

To estimate growth rates for each species, three replicate bottles of soil extract medium were inoculated with 15-20 mL of dense *Loxodes* culture, and filled to 100 mL with soil extract medium. Samples were taken as above for counting twice weekly, and fed with *Chlamydomonas* as described above after counting. After cultures appeared to have reached saturation density, they were transferred to new bottles, and new soil extract medium was added to a total volume of 200 mL each, and monitored for another week. Cell count data are available at doi:10.17617/3.4ZQNBK.

### Harvesting of cells from bulk culture for experiments

Cultures were filtered through a layer of cotton gauze swabs (Paul Hartmann AG) to remove flocculent debris, pouring gently to avoid breaking up debris into smaller pieces that can pass through the gauze. Ciliates were concentrated by centrifugation (*Loxodes magnus* 120 rcf, *L. striatus* 240 rcf; 1 min; room temperature) in pear-shaped flasks using an oil-testing centrifuge (Rotanta 460, Hettich, Ref 5650), as previously described for the ciliate *Paramecium* (Beisson et al., 2010b; Sonneborn, 1950), and then resuspended in a smaller volume of new SMB medium.

### Preparation of fixatives

ZFAE solution is a modification of Nissenbaum’s fixative (Nissenbaum, 1953), and consists of the following components mixed together shortly before use: 10 volumes of 0.25 M zinc sulfate (Sigma-Aldrich), 2 volumes of 36.5-38% (w/v) formaldehyde solution (Sigma-Aldrich), 2 volumes of glacial acetic acid (Roth), 5 volumes absolute ethanol (Roth). Commercial formalin (37% formaldehyde, 10% methanol as stabilizer) has also been used instead of formaldehyde without any apparent difference in results.

Other fixatives tested were 4% formaldehyde (w/v) in phosphate buffered saline (FA-PBS), 4% formaldehyde (w/v) in SMB medium (FA-SMB), and ice-cold methanol (MeOH).

### Fixation of ciliate cells

For ZFAE fixative, cells were fixed with twice the volume of fixative solution for 1 h at room temperature (RT), centrifuged (100 rcf; 2 min; RT), and resuspended in SMB. For FA-SMB and FA-PBS, 30 µL of concentrated cells were added to 1 mL of fixative solution, fixed for 1 h at RT, centrifuged, and resuspended in respective buffer solution (SMB or PBS). For MeOH, 20 µL of concentrated cells were added to 1 mL ice-cold MeOH, fixed for 5 min, centrifuged, and resuspended in SMB. ZFAE-fixed cells can also be resuspended again in 70% ethanol and stored at 4 ºC for later use.

Live and fixed cells were imaged with differential interference contrast (DIC) on a Nikon Eclipse Ts2R inverted microscope with either 20× or 40× objective and Zeiss AxioCam HRC camera. Contrast and brightness were adjusted in the Zeiss AxioVision software to achieve a similar background brightness.

### Immunofluorescence

This protocol was adapted from (Beisson et al., 2010c). Primary antibody against alpha-tubulin (raised in rat, Abcam ab6161) was diluted 1:100 (v/v) in 3% (w/v) bovine serum albumin (BSA, Sigma-Aldrich) in TBSTEM buffer (10 mM EGTA, 2 mM MgCl_2_, 150 mM NaCl, 10 mM Tris, 1% Tween-20, pH 7.4). Secondary antibody (goat anti-rat, Alexa Fluor 488 labeled, Abcam ab150157) was diluted 1:200 (v/v) in 3% BSA / TBSTEM. DAPI was diluted to 1 µg/mL in 3% BSA / TBSTEM. All resuspension steps were performed by first centrifuging (1000 rcf; 1 min; room temperature), then removing supernatant, and resuspending the pellet by pipetting up and down. Fixed cells were resuspended in 0.5 mL diluted primary antibody and incubated for 10-60 min at room temperature (RT). Cells were washed for 5-10 min RT in 1.5 mL of 3% BSA / TBSTEM, then incubated for 10-30 min RT in 0.5 mL diluted secondary antibody. Cells were counterstained for >5 min at RT in 1 mL DAPI / BSA, then resuspended in 15 µL ProLong Gold (P36930, Thermo Fisher), mixed by gentle stirring, and mounted under a coverslip. Preparations were cured overnight at RT before imaging. Coverslips can be sealed with nail polish. Sealed slides kept at 4 °C for several months can still be used for imaging. DIC and fluorescence images were imaged on a Zeiss AxioImager Z.1 with 40× objective, with Zeiss Filter Set 49 for DAPI and Zeiss Filter Set 38HE for Alexa Fluor 488, and Zeiss AxioCam HRc Rev.3 camera.

### Safe disposal of ZFAE fixative

Excess sodium bicarbonate was added to neutralize the acetic acid and to precipitate zinc as zinc bicarbonate, which was filtered out and disposed of as solid chemical waste to minimize waste volume. Formaldehyde in the filtrate was inactivated by adding excess glycine or milk powder and stirring for >1 h.

## Results

### Growth of Loxodes magnus in liquid soil medium

*Loxodes* cultures were maintained in soil-liquid test tubes as well as in liquid soil extract medium. The media were visibly turbid a few days after inoculation, because neither cultivation medium was axenic, and both depend on the respiration of bacteria co-isolated with *Loxodes* to produce the micro-oxic conditions preferred by the ciliates.

Growth rates of *Loxodes* were estimated in liquid culture, because in soil-liquid tubes many cells also reside in the soil and cannot be easily counted. *L. striatus* cell counts did not show much change for the first 6 days, then entered an exponential growth phase (linear region on the log-scaled plot) (Figure 2A, C). This continued after the culture was expanded by adding fresh medium, but one replicate experienced a crash in cell density just before the second expansion. The experiment did not continue long enough to observe a sustained plateau in cell density, so the saturation density of *L. striatus* is probably above the highest density observed of ca. 2^10^ ≈ 1000 cells / mL. In comparison, *L. magnus* showed exponential growth for the first 10 days; subsequently cell density appeared to plateau at about 2^9^ ≈ 500 cells / mL (Figure 2B, D). Cell density soon returned to this saturation value after each time that new medium was added. Doubling times, as judged from the linear regions of the log-scaled density plot, were about 7-9 days for *L. striatus*, vs. 4-8 days for *L. magnus*, although *L. striatus* is a smaller species and was hence expected to divide more rapidly. We hypothesize that the actual division rate of individual *L. striatus* cells was indeed faster than *L. magnus*, but that the overall doubling rate was limited by the available algal food, which was supplied at the same rate for both strains.

**Figure 2.**
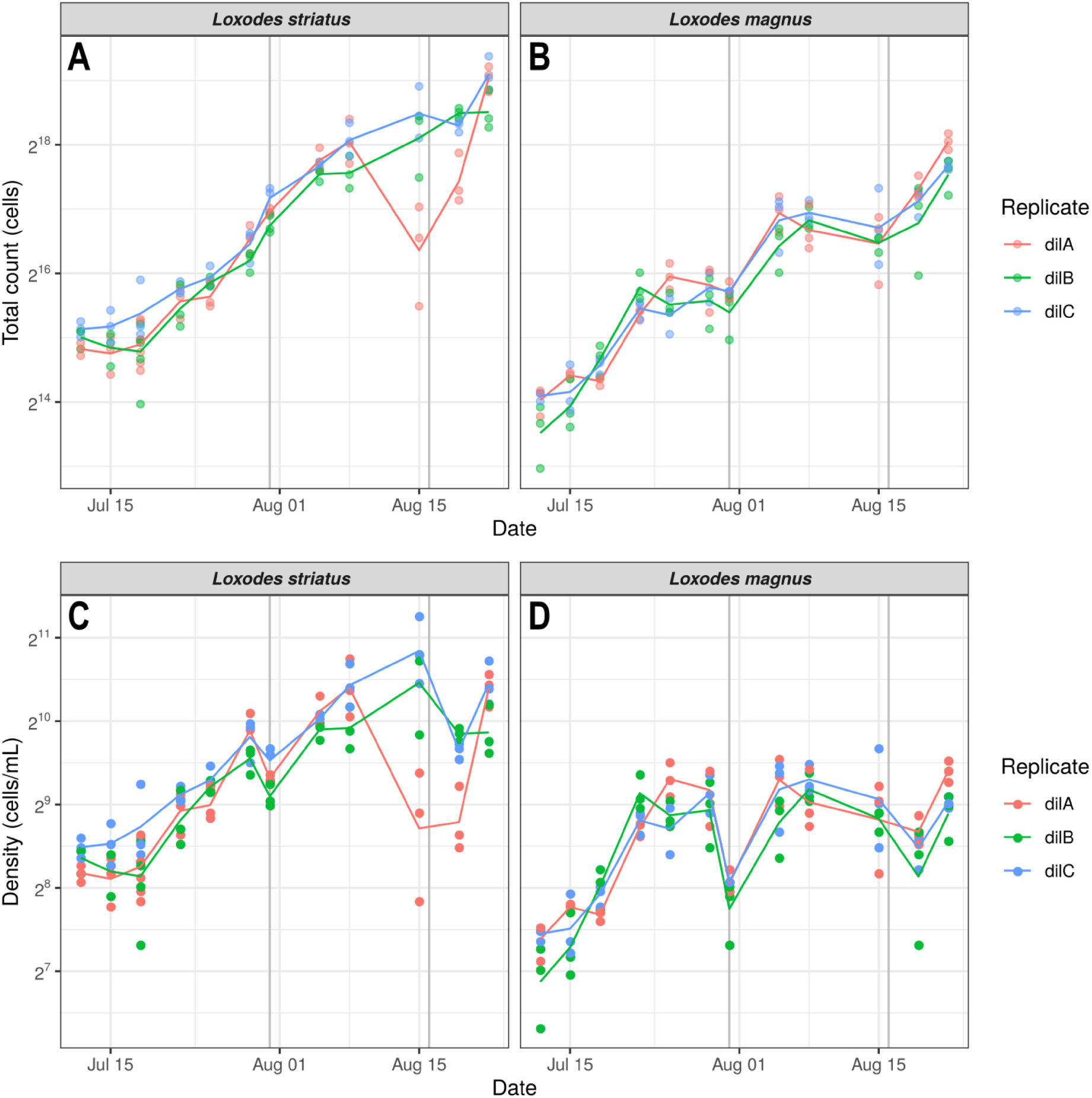
Growth of *Loxodes striatus* (A, C) and *L. magnus* (B, D) in soil extract medium, each strain grown in triplicate (colors), plotted as total counts (above) vs. cell density (below). Points represent each aliquot taken for counting (3 per replicate), lines are mean counts per replicate. Dark grey vertical lines mark dates when total culture volume was expanded two-fold by adding fresh medium.

### Comparison of fixation by ZFAE vs. formaldehyde or methanol

We first evaluated the effectiveness of formaldehyde and methanol, fixatives typically used in immunofluorescence protocols, against *Loxodes magnus* and the heterotrich *Blepharisma stoltei*, which is also known to be challenging to fix. Formaldehyde solutions were buffered with either PBS or SMB, because live cells could be resuspended in these solutions without lysing or losing motility.

Neither 4% formaldehyde in phosphate buffered saline (FA-PBS), 4% formaldehyde in SMB medium (FA-SMB), nor ice-cold methanol (MeOH) could fix *Loxodes* cells effectively for downstream applications. *Loxodes* cells fixed with these fixatives were swollen and soft, and most cells partly disintegrated with the handling required to mount cells under a coverslip, although nuclei appeared to be intact (Figure 3A-D). *Blepharisma* cells fixed with FA-SMB or FA-PBS were also swollen compared to live morphology, although more cells remained intact (Figure 3E-G). MeOH fixation caused the pellicles of *Blepharisma* cells to swell and partially detach from the rest of the cytoplasm, but adoral membranelles were less disaggregated compared to FA-PBS and FA-SMB (Figure 3H).

**Figure 3.**
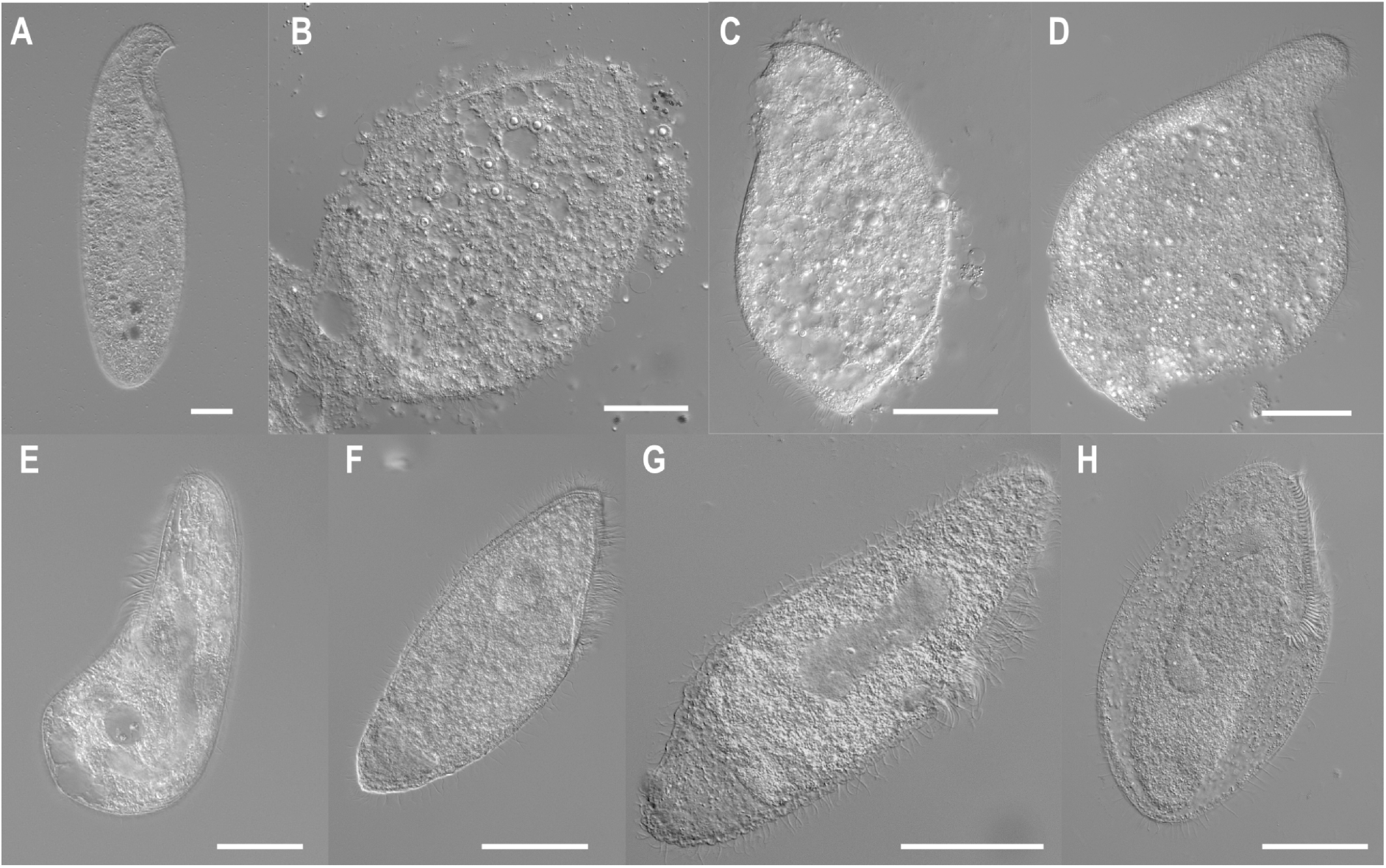
Comparison of live cells vs. fixed with conventional fixatives, *Loxodes magnus* (above) and *Blepharisma stoltei* (below), imaged with differential interference contrast. Live cells (A,E); 4% formaldehyde/SMB (B,F); 4% formaldehyde/PBS (C,G); ice-cold methanol (D,H). All scale bars: 50 µm.

In comparison, ZFAE-fixed cells, both for *Loxodes* and *8/epharisma*, remained intact and did not disintegrate on handling (Figure 4A, D). Cells shrank relative to live material but largely kept their original shape, although cilia were stubby or lost. Cell pigments (pink for *8/epharisma*, yellow for *Loxodes)* were also partially extracted by ZFAE, probably because of the ethanol in the fixative. For *8/epharisma*, the posterior contractile vacuole was clearly visible in ZFAE-fixed cells, unlike with other fixatives. However the Muller vesicles of *Loxodes* were not observed in fixed samples, including with ZFAE.

**Figure 4.**
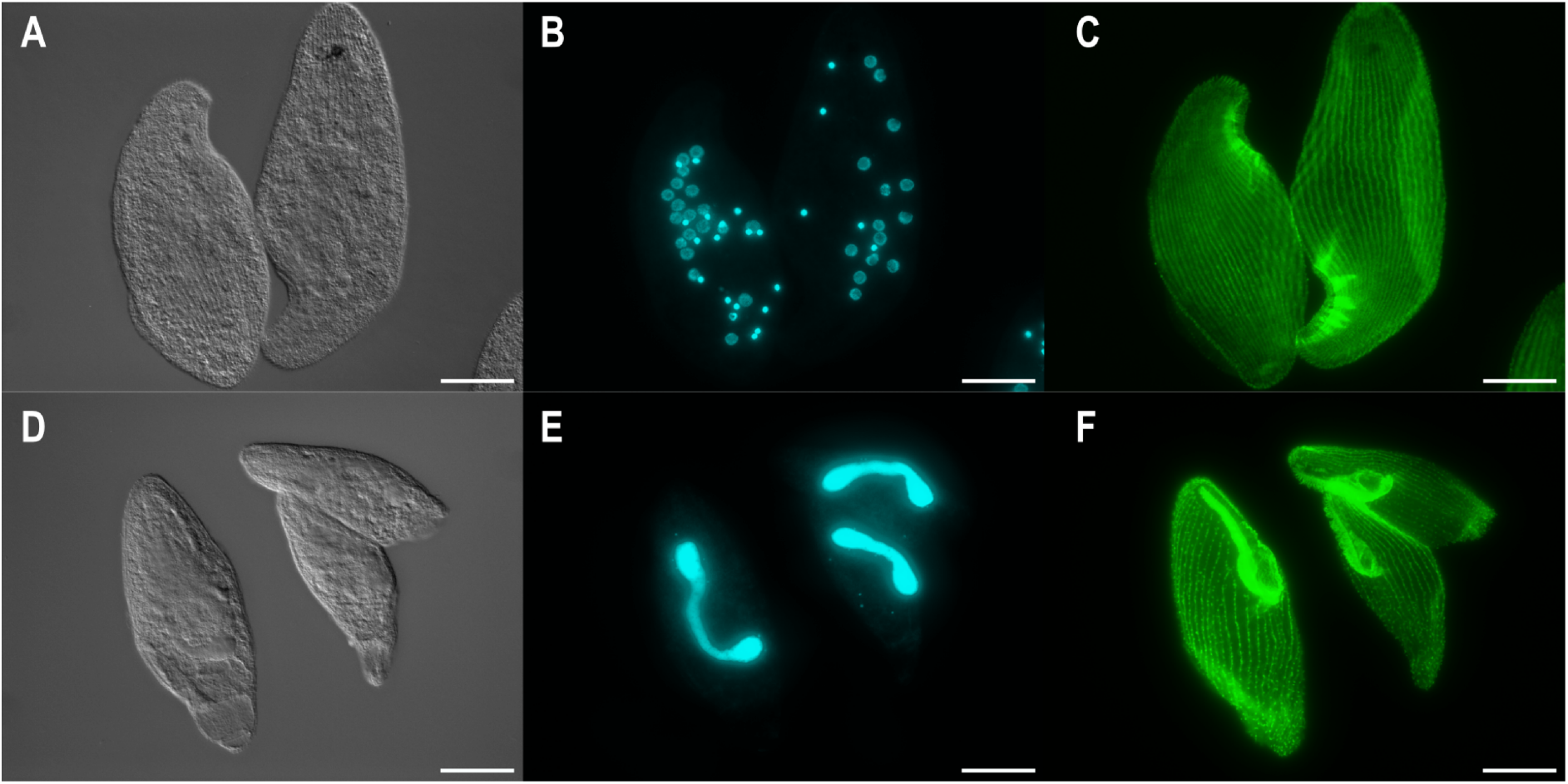
*Loxodes magnus* (above) and *Blepharisma stoltei* (below) cells fixed with ZFAE. Morphology imaged with DIC (left), and false color fluorescence micrographs of nuclei labeled with DAPI (center), and alpha-tubulin labeled with secondary immunofluorescence (right). All scale bars: 50 µm.

### Compatibility of ZFAE fixation with immunofluorescence

Ciliate cells fixed with ZFAE were compatible with indirect immunofluorescence and DAPI staining of nuclear DNA.

Indirect immunofluorescence with a primary antibody against alpha-tubulin labeled linear structures in *Loxodes magnus* that corresponded with the expected appearance and number of somatic kineties (Figure 4C). Cells were laterally flattened, with most ciliation on the right side (ca. 25 somatic kineties), vs. two somatic kineties on the left side that were more faintly labeled. Oral kineties were more difficult to distinguish because bundles of fibrous structures were brightly labeled in the oral region and obscured the signal from other structures. These bundles appear to be nematodesmata associated with the paroral kineties and the left pseudobuccal kinety (Foissner and Rieder, 1983; Klindworth and Bardele, 1996). The vestibular extension (“pharynx”) was also labeled. Nuclei stained with DAPI had their characteristic morphology: micronuclei were densely and uniformly stained, whereas macronuclei were less intensely stained, and the central nucleolus was only weakly labeled by DAPI (Figure 4B).

*Blepharisma stoltei* cells were spindle-shaped and ciliated all around, in line with previous morphological descriptions (Giese, 1973). Similarly to *Loxodes*, the oral region, particularly the adoral zone of membranelles, was brightly labeled so details of the oral ciliature were difficult to distinguish when overlapping (Figure 4F). The elongate macronucleus was strongly labeled with DAPI, and small spherical micronuclei were also observed beside the macronuclei (Figure 4E).

## Discussion

We have shown that *Loxodes* ciliates can be grown in bulk in liquid medium, and that the new ZFAE fixative can be used to fix fragile ciliates like *Loxodes* and *Blepharisma* for fluorescence microscopy applications.

Cultivation of *Loxodes* and indeed most ciliates is xenic, i.e. ambient bacteria are present, but the target ciliate and food algae should be the only eukaryotes present. *Loxodes* cultures often collapsed or died out some time after the initial isolation, which was likely influenced by the composition or dynamics of the accompanying bacterial community. Therefore, multiple isolates should be maintained and subcultured for some time, before the most consistently growing ones are chosen for expansion in bulk liquid culture. In our laboratory, the most stable strains have been maintained for over two years in both soil-liquid tubes and soil extract medium.

Successful cultivation of *Loxodes* appears to require soil or soil extract, because it is a component of most reported protocols (Supplementary Information). As the composition of soil can vary greatly, it is necessary to test different local soils when setting up a culture, starting with soil from the site where the ciliates were collected. Some previous studies have reported bubbling liquid culture medium with nitrogen gas, or using a chamber flushed with nitrogen, to achieve low oxygen concentrations (Buonanno et al., 2005; Nagel et al., 1997). We did not find this necessary, presumably because the soil extract contains sufficient organic carbon (195 mg / L total organic carbon) to support bacterial growth that is fast enough to consume excess oxygen, and diffusion of oxygen is limited by using bottles or tubes vs. shallow dishes. Indeed, *Loxodes* cultures tend to die out if bottles were completely filled without leaving any headspace, consistent with previous reports that *Loxodes* cannot tolerate complete anoxia (Goulder, 1980). Optimal growth temperatures may be strain and locality dependent. Our cultures are maintained at room temperature (24 ºC), whereas attempts to move backup cultures to lower temperatures (18 ºC, 4 ºC) were not successful. *Loxodes* also appears to be flexible in their choice of food algae. Both *L. striatus* and *L. magnus* can also be fed with *Euglena gracilis* and *Chlorogonium* sp., but *Chlamydomonas* was chosen because it was the easiest to grow.

The ZFAE fixative fills a useful niche in the ciliatology toolkit, as a fixative that works for at least two known “difficult” taxa, and which is also suitable for fluorescence microscopy, which opens the door to applying cell biology methods to a wider range of ciliate model species. Fluorescent labeling of the cytoskeleton has also been suggested as an alternative method for species identification and description in ciliates vs. traditional silver staining methods (Hirst et al., 2011; Trogant et al., 2020), because the reagents and expertise are more readily available from commercial suppliers, and because they can potentially be imaged at higher resolution, e.g. with confocal microscopy. Methanol has been used to fix *Blepharisma* for immunofluorescence (Santangelo and Bruno, 2001), but the reported protocol involves immobilizing cells in a thin film of culture medium on a depression slide before adding fixative. Similarly, a protocol used to fix *Loxodes* for DAPI staining adheres them to glass slides (Munyenyembe et al., 2021). These methods require careful manual manipulation and limit the numbers of cells that can be fixed. In contrast, ZFAE is more suitable for large numbers of cells, and is simpler as all steps can be performed at room temperature in microcentrifuge tubes. Drawbacks of ZFAE, however, are that surface ciliation is poorly preserved, and morphology may be distorted because of shrinkage during fixation.

The key difference between ZFAE and the original Nissenbaum’s fixative is the substitution of mercuric chloride with zinc sulfate. Mercuric salts were historically widely used in histology (Hopwood, 1972), but are rarely used today because of their toxicity and the expense of chemical disposal (Layton et al., 2019). Zinc salts have successfully substituted for mercuric salts in several applications (Garcia et al., 1993; Lykidis et al., 2007), and are thought to work via a similar mechanism. The original Nissenbaum’s fixative was used for simultaneous fixation and enrobement, i.e. adhesion of specimens to glass slides. In contrast, ZFAE fixative is not effective at enrobement, so fixation in tubes or vials is recommended. Nonetheless, this is an advantage for protocols like immunofluorescence, and it is also more convenient to store fixed samples in tubes for later use.

Finally, ZFAE is composed of inexpensive chemicals, and the disposal of fixative waste is more convenient and safer than for mercuric chloride-containing fixatives. Besides protecting laboratory workers, this also saves on shipping, storage, and disposal costs. Alternatively, 40% glyoxal solution can be used in place of formalin as a less toxic substitute (Richter et al., 2018). Standard precautions (fume hood, personal protective equipment) should still be taken.

## Acknowledgements

This work pays tribute to the contributions of Bland Finlay to our knowledge of *Loxodes* and the microbial world. We thank T. Harumoto and M. Sugiura for providing the *Blepharisma* culture, K. Eisler for the gift of *Loxodes* cultures from the former teaching collection of the University of Tübingen, S. Mattes for routine maintenance of our cultures, B. Seah Mikitish for translation from Russian, and G. Esteban for the invitation to contribute this article.

