## Supplementary Information for "Improved methods for bulk cultivation and fixation of *Loxodes* ciliates for fluorescence microscopy"

Cultivation methods for *Loxodes* ciliates reported in published papers or from personal communication. All methods require either actual soil or soil extract, which is not chemically defined but whose composition is highly variable.

### Nagel method

- Species: *Loxodes striatus* (strain originally provided by B. Finlay)
- References: Nagel et al., 1997; Neugebauer et al., 1998
- Medium:
  - Mineral salts medium: 0.5 mM KCl, 1 mM $CaCl_2$, 0.1 mM $MgSO_4$, 0.41 mM $Na_2HPO_4$, 0.18 mM $NaH_2PO_4$, (phosphate buffer yielding final pH 7.0), deoxygenated by gassing with $N_2$.
  - Alternative: Soil solution (composition not specified)
- Method:
  - Add one (unpolished) rice grain and 20 mL medium per test tube
  - Keep at 22 °C or 18 °C in darkness
  - Medium is presumably bacterized because of the rice grain
- Feeding: *Euglena gracilis*, twice weekly
- Notes:
  - No apparent difference in size or behavior of Loxodes grown in mineral salts vs. soil media
  - Cells were starved 2-5 days before experiments

### Buonanno method

- Species: *Loxodes striatus* strain CH1, isolated from Chieti, Italy.
- Reference: Buonanno et al., 2005
- Medium: Soil extract medium diluted with SMB-III. Ratio not specified.
  - SMB-III medium: 1.5 mM NaCl, 0.05 mM KCl, 0.4 mM $CaCl_2$, 0.05 mM $MgCl_2$, 0.05 mM $MgSO_4$, 2 mM Na-phosphate buffer, final pH 6.8, $2 \times 10^{-3}$ mM EDTA
- Method:
  - Fill beakers with 3-5 mm of autoclaved soil and 80-120 mL SMB-III medium, kept in a chamber under $N_2$ gas in the dark, gas is exchanged daily.
- Feeding: *Chlorogonium elongatum*, every 2 days

### Bobyleva method

- Species: Various *Loxodes* spp.

- Reference: Bobyleva, 1980
- Method:
  - Fill garden soil to 3 cm in test tubes, add water to 8-10 cm, autoclave, store at 4 °C before use.
  - Soil extract used for topping up test tubes or for incubating clones: 300 g soil in 1.5 L water, autoclave, filter, and re-autoclave. Store at 4 °C.
- Feeding:
  - Cultures at 10 °C: 1 mL *Euglena* culture, twice weekly subcultured once every two months.
  - Cultures at 17 °C: 1 mL *Euglena* culture, three times weekly, subcultured once every month
  - Reserve cultures at 10 °C: 1 mL *Euglena* culture once a week, kept for up to 4-6 months

### Fenchel & Finlay method

- Species: *Loxodes striatus* and *L. magnus*
- Reference: Fenchel and Finlay, 1984
- Method:
  - Soil extract medium (composition not specified) completely filled in tissue culture flask. Not leaving a headspace presumably reduces $O_2$ concentration.
  - Alternative: Test tubes with 1-2 cm autoclaved soil and soil extract medium
  - Cultures kept at 10, 15, or 20 °C
- Notes: Bacteria in the medium lower the $O_2$ concentrations by respiration
- Feeding: *Euglena gracilis*

### Tübingen method (unpublished)

- Species: *Loxodes striatus* and *L. magnus*
- Reference: Unpublished protocol from former teaching culture collection at the University of Tübingen, courtesy of K. Eisler
- Method:
  - Fill test tubes (about 20 cm long) 1/5 to 1/3 with air-dried, light loamy garden soil (without fertilizer), and top up to the rim with boiled, cooled Volvic water
  - On the next day: Inoculate with 2-3 pipettefuls of a dense *Loxodes* culture
  - Maintain at 15 °C
- Feeding: Pipetteful of *Euglena*, once weekly
